# Draft genome sequence of a popular Indian tea genotype TV-1 [*Camellia assamica L*. (*O*). Kunze]

**DOI:** 10.1101/762161

**Authors:** Tapan Kumar Mondal, Hukam Chand Rawal, Biswajit Bera, P. Mohan Kumar, Mrityunjay Choubey, Gargi Saha, Budhadeb Das, Tanoy Bandyopadhyay, R. Victor J. Ilango, Tilak Raj Sharma, Anup Barua, B. Radhakrishnan, Nagendra Kumar Singh

**Affiliations:** ICAR-National Institute for Plant Biotechnology, LBS Centre, IARI, New Delhi, India; Tea Board, Ministry of Commerce and Industry, Govt. of India, 14, B.T.M. Sarani, Kolkata, 700 001, India; Tocklai Tea Research Institute, Tea Research Association, Jorhat, Assam, 785008, India; UPASI Tea Research Foundation, Tea Research Institute, Coimbatore, Tamil Nadu, India

**Keywords:** Camellia, *C. assamica*, Genome size, Next generation sequencing, Tea, Whole genome sequence, assembly, Woody plant

## Abstract

*Camellia assamica L*. (*O*). Kuntze. var. TV-1 from Assam, India is one of the most important tea cultivar of Indian Tea Industry which botanically belongs to Assam type and has distinct morphological as well as chemical constituent in leaf than China type of tea. We here present the first draft genome sequence of TV-1, assembled in 2.93 Gb size comprising of 14,824 scaffolds and covering 97.66% of the predicted genome size (3.0 Gb) as determined by flow cytometry. The N50 value, longest and average contig size for this draft genome was found to be as 538.96 Kb, 3.64 Mb and 197.99 Kb, respectively. There were 495,747 SSRs (Di- to Hexa-mers) in the genome and the repeat contents accounted for 71.87% of the genome. About 94.40% genome completeness was predicted with BUSCO, maximum among the published and reported tea genomes. The clustering of the assembled genome with Hi-C data resulted in 15 clusters ranging from the size of 631.46 Mb to 14.39 Mb and accounting 99.32% of assembled genome. This data will act as the starting point for unravelling the important genes and differentiating Indian tea from other commercially available tea varieties.

## Introduction

Tea is a popular non-alcoholic beverage of the world. It is an industrial crop that provides ecofriendly environment and generate large number of employment particularly in the backward places of developing countries such as India, China and other African countries. However being woody perennial plant, varietal improvement is very slow in tea (Mondal et al., 2004) and hence application of biotechnology tools such as transgenic, genetic manipulation via CRISPR/CAS-9 and marker assisted breeding will be better alternative approach. However one of the prerequisite for applying such approaches is the requirement to decode the genome to speed up the genetic improvement of tea.

Although multiple efforts has been made for decoding the genome of China type tea with two different China type varieties, *C. sinensis* var. a*ssamica* (CSA) (Xia et al. 2017) and *C. sinensis* var. *sinensis* (CSS) (Wei et al. 2018, Xia et al. 2019) data but no attempt has been reported so far for decoding the Indian tea (*C. assamica*) genome. It is noteworthy to mention here that Indian tea is mainly dominated by Assam type of tea which is genetically and morphologically distinct from China type. Therefore to understand the genetic make-up of the Indian tea, we have made an attempt to decode the TV-1 genotype using NGS data including Illumina, PacBio and Hi-C long reads. TV-1 is the oldest clone that was released in 1949 and covers a significant area under cultivation in Assam, India. Besides, it is used as a parent of several tea breeding programs in India (Deka et al., 2006). In addition, it has low heterozygosity among the major popular tea cultivars that are used in Indian Tea Industry (unpublished data).

## Materials and methods

### Genome sequencing

The flow cytometer technique (Dolezel et al., 2007) was used for genome size estimation of TV-1 by using *Oryza sativa* and *Pisum sativum* as reference standard genome. The tissue of 2^nd^ young leaf was collected from the tea genotype, TV-1 from Tocklai Tea Research Institute, Assam, India for the isolation of genomic DNA using the CTAB protocol (Doyle and Doyle 1987) to prepare various sequencing libraries such as Illumina, PacBio and Hi-C library.Quality of the DNA was checked by a NanoDrop D-1000 spectrophotometer (NanoDrop Technologies, Wilmington, DE) and Qubit Fluorometer. The short Paired-end (PE) of 151 bp long reads and three different Mate-pair (MP) libraries of 2.5 -4 kb, 4-6 kb, 8-11 kb size were made following the standard Illumina protocols (Illumina, San Diego, CA) and sequenced with HiSeq4000 platform (Illumina, San Diego, CA). For preparation of PacBio SMRT bell libraries 3-5 μg high-quality, high molecular weight genomic DNA was used. The integrity of input material was checked on pulsed field gel electrophoresis and DNA sample of predominantly >50 kb was used for production of six large insert (7-20 kb) libraries by sequencing with PacBio Sequel.DNA for the Hi-C experiment was prepared at Phase Genomics after incubating at 27 °C for 30 minutes with periodic mixing by inversion followed by addition of Glycine (final concentration of 0.1 g/10 mL) to quench crosslinking, an additional incubation at 27 °C for 20 minutes with periodic inversion, and removal of supernatant. The samples were then sequenced by HiSeq4000 platform (Illumina, San Diego, CA).

### Genome assembly

The high-quality PacBio reads were subjected to Platanus (Kajitani et al. 2014) for first level PacBio assembly followed by error correction and scaffolding by Npscarf v1.7 (Cao et al., 2017) using high-quality (Q-30) processed and adaptor trimmed Illumina PE and MP reads. GapCloser (Luo et al., 2012) and Sealer (Paulino et al., 2015) were used with additional Illumina PE reads to remove gaps and obtain final assembly. Finally the clustering was performed with Hi-C data using LACHESIS (Burton et al. 2013) to assign assembled scaffolds into different clusters.

### Assessment of Genome Completeness

The genome completeness was evaluated initially with traditional Core Eukaryotic Genes Mapping Approach (CEGMA) tool for the presence of 248 ultra-conserved CEGs in the assembled genome (Parra et al., 2007). Further Benchmarking Universal Single-Copy Orthologs (BUSCO) tool was used for the final assessment of genome completeness with 1375 gene sets from Embryophyta (odb10) lineages (Simao et al., 2015).

### Determination of repetitive content

Repeat Modeler (Smit and Hubley, 2008) was used to identify the *de novo* repeat families and mask the total repetitive content in the assembled genome. For detecting the frequency of microsatellites or Simple Sequence Repeats (SSRs) in the genome, MIcroSAtellite identification tool (MISA) was used with motif size ranging from di- to hexa-nucleotide (Thiel et al. 2003).

### Assembly of organelle genomes

Using the PacBio and Illumina MP reads, we have assembled the organelle genomes of *C. assamica* (Rawal et al. 2019). We assembled the mitochondrial and chloroplast genome of TV-1 with 707,441bp and 157,353 bp length, respectively. While this was the first and only effort to decode the mitochondrial genome of any *Camelia* species, the chloroplast genome is the first one for *C. assamica*. The phylogenetic analysis based on these organelle genomes has shown that *C. assamica* was closely related to *C. sinensis* and *C. leptophylla*. The generation of nuclear genome will add up in understanding the evolution and origin of *C. assamica* at genomic and gene level. It will further provide data to identify and understand the genetic variations among Indian and China type tea.

## Results and discussion

### Genome size and generation of raw data

We report the draft genome assembly of a popular Indian tea clone TV-1. The genome size of this genotype was estimated to be 3.0 Gb with flow cytometer technique which is a close proximity with other tea genome size (Huang et al., 2013). Considering this 3.0 Gb genome size, about 136X and 13X of raw data were generated with Illumina (including PE and MP reads) and PacBio reads, respectively to achieve a total sequencing depth of 149.29X (Table 1). For clustering the assembled scaffolds, a total of 147.17 Gb of Hi-C reads, accounting 49.06X depth, were generated.

**Table 1:** Statistics of TV-1 genome assembly and its comparison with published tea genomes

| Attributes | *C. assamica*<br>(Indian tea)<br>(TV-1) | *C. sinensis* var.<br>*sinensis*<br>(Wei et al. 2018;<br>Xia et al. 2019) | *C. sinensis* var.<br>*assamica*<br>(Xia et al. 2017) |
| --- | --- | --- | --- |
| **Assembly** |  |  |  |
| Genome-sequencing Depth (X) | 149.29 | 739.79 | 159.43 |
| Estimated Genome Size (Gb) | 3.00 | 3.08 | 3.09 |
| Assembled Genome Size (Gb) | 2.93 | 3.14 | 3.02 |
| Predicted Coverage of Assembled Genome (%) | 97.66 | 95.07 | 97.73 |
| Number of Scaffolds | 14,824 | 14,051 | 37,618 |
| Total Length of Scaffolds (bp) | 2,934,935,288 | 3,141,536,798 | 3,021,230,785 |
| Longest Scaffold (bp) | 3,635,958 | 7,310,916 | 3,505,831 |
| Smallest Scaffold (bp) | 1000 | 5000 | 500 |
| Average Scaffold Length (bp) | 197,985.3 | 223,581.01 | 80,313.43 |
| GC % | 39.57 | 37.84 | 42.31 |
| Non-ATGC Characters | 359,701,292 | 247,754,689 | 446,294,562 |
| Non-ATGC % | 12.25 | 7.89 | 16.89 |
| Scaffolds >= 1 Mbp | 462 | 1,102 | 354 |
| N50 of Scaffolds (bp) | 538,958 | 1,397,810 | 449,457 |
| **Repeat Analysis** |  |  |  |
| Masked Repeat Sequence Length (bp) | 1,853,117,835 | 1,861,774,995 | 1,753,247,986 |
| Repeat Sequences in the genome (%) | 71.87 | 64.42 | 80.89 |
| **Genome Completeness** |  |  |  |
| CEGMA: Complete % | 76.61 | 76.61 | 76.21 |
| Partial % | 95.97 | 93.95 | 94.35 |
| BUSCO: Complete % | 94.40 | 92.40 | 85.50 |
| Fragmented % | 2.00 | 3.10 | 2.60 |
| Missing % | 3.60 | 4.50 | 11.90 |
| **Hi-C Clustering** |  |  |  |
| Clusters | 15 (2.91 Gb) | - | - |
| Un-clustered sequences | 393 (22.35 Mb) | - | - |
| N50 | 419,173,006 | - | - |
| Largest Cluster size (bp) | 631,461,571 | - | - |

### Genome assembly and repeat analysis

The assembly resulted in a draft genome of 2.93 Gb covering 97.66% of estimated genome size with N50 size of 538,958 bp (Table 1). There were 14,824 scaffolds (>1kb) in the assembly with the largest contig was of 3,635,958 bp size, and comprising of 82.92% scaffolds >10 kb and 462 scaffolds >1 Mb size. There was only 12.25% non-ATGC or gaps in the assembled genome and the repetitive content accounted for 71.87% of the genome. A total of 495,747 SSRs were identified with MISA comprising 78.57%, 15.39%, 3.81%, 1.16%, and 1.07% for Di-, Tri-, Tetra-, Penta- and Hexa-nucleotide repeats (Table 2).

**Table 2:** Number of different types SSRs in the *C. assamica* genome

| S. No. | Motif-type | Minimum Repeat number | Number of SSRs | % of SSRs |
| --- | --- | --- | --- | --- |
| 1. | Di | 6 | 389523 | 78.57 |
| 2. | Tri | 5 | 76290 | 15.39 |
| 3. | Tetra | 5 | 18895 | 3.81 |
| 4. | Penta | 5 | 5745 | 1.16 |
| 5. | Hexa | 5 | 5294 | 1.07 |

Hi-C data clustered the 99.32% (2.93 Gb) of the assembled genome into 15 clusters (equivalent to chromosome number in the tea) with N50 size of more than 419 Mb (Figure 1). Eight of these 15 clusters were found to be more than 100 Mb in size with the largest cluster of 631.46 Mb length. Hi-C served the purpose of clustering with the un-clustered data accounted for just 22.35 Mb, which is less than 1% of the assembled genome.

**Figure 1:**
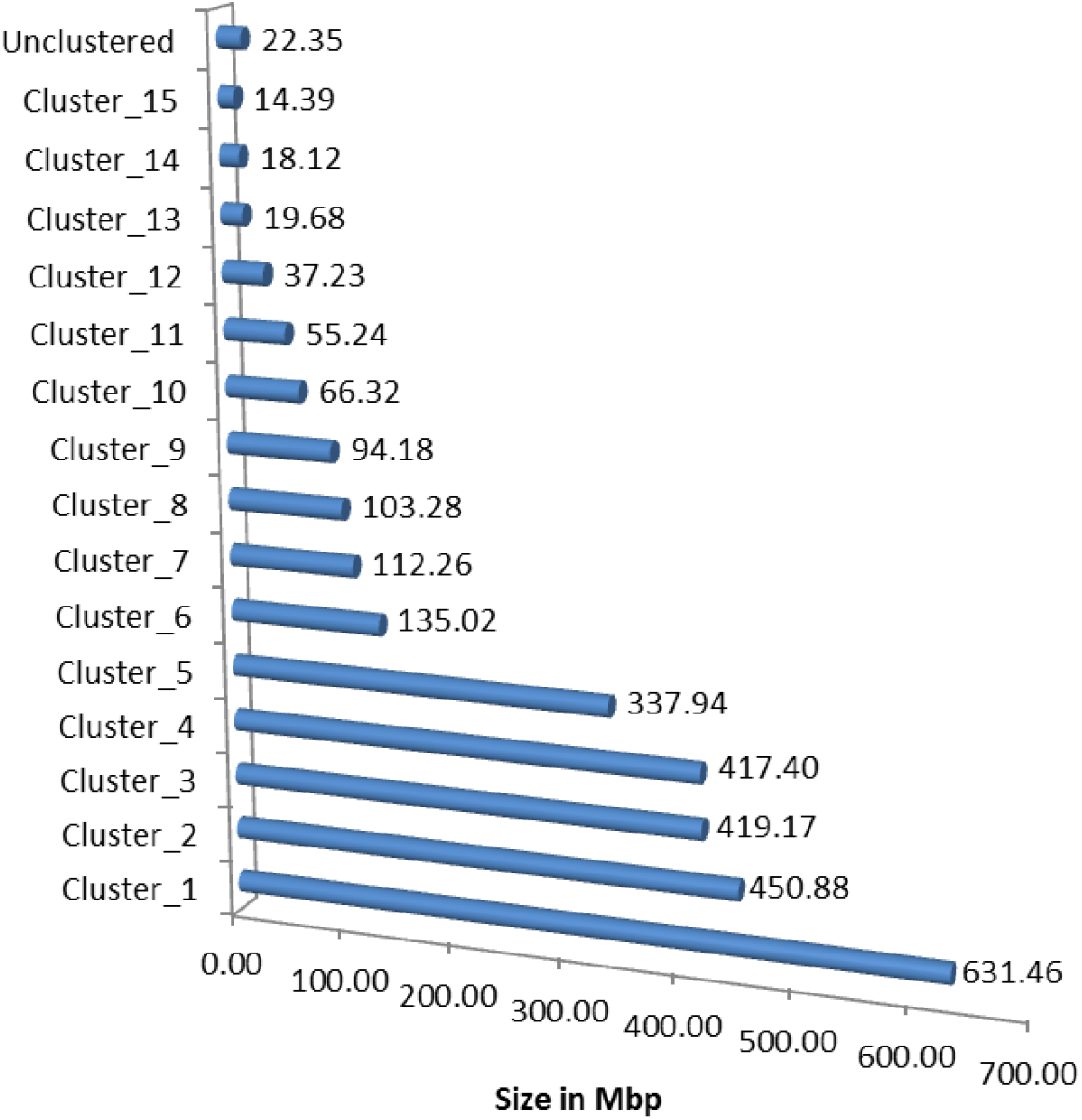
Size distribution of clusters made with HI-C data

### Completeness check

The genome completeness (94.40%) was evaluated with BUSCO (Simao et al. 2015) and only 3.60% of the conserved core eukaryotic genes were missing in the assembly while 2.00% were present as fragmented, indicating the better completeness as compared with the other two published tea genomes (Xia et al. 2017; Wei et al. 2018, Xia et al. 2019). Similarly, the completeness check with CEGMA indicated that 76.61% complete and 95.97% partial/complete conserved eukaryotic genes were present in the present assembly, comparatively as good as that was in the published ones.

### Comparative analysis

The advancement in NGS technology aids well in our achievement by generating reference level assembly with less than 150X coverage, as compared to the previous assemblies of tea genomes with upto 740X sequencing depth. While genome completeness percentage was found better than both of the published tea genomes, most of the aspects of our genome assembly were better than CSA assembly which included number of scaffolds, longest and smallest scaffold size, non-atgc or gap contents and more importantly the N50 size (Table 1). Comparing to CSS assembly, although we need to work on reducing the gaps which could also aids in increasing the N50 size, otherwise we achieve better or almost similar aspects including BUSCO completeness score (94.40% vs 92.40%), CEGMA score (76.61% each for complete and 95.97 vs 93.95% for partial genes), Genome coverage (97.66% vs 95.07%), Scaffolds (14,390 vs 14,051 with ≥5000 bp length). Moreover, as mentioned above, this is the first cluster level assembly with over 99% data clustered into 15 clusters.

### Future aspects

The availability of the *C. assamica* genome sequence is a first step towards unravelling the genetic variations among Indian and China type tea, and studying the evolutionary origin of *C. assamica* at genomic and gene level. This will also serve as the baseline data to identify the unique genetic variation of Indian tea and will aid to identify the agronomically important genes which are useful for Indian tea Industry. Besides, this data will be useful for several other works related to genetic improvement of Indian tea.

## FUNDING

The work was funded by National Tea Research Foundation, Tea Board, Ministry of Commerce, Kolkata, West Bengal, India

## ACKNOWLEDGMENTS

The authors are grateful to Mr. Abhishek Mazumder for his help to get the sample, Miss Megha Rohila for conducting the flow cytometer experiments. Miss Deepti Varshney, Dr. Priyanka Jain, Dr. Himanshu Dubey and Dr. P. Mahadani for their technical help in some *in silico* analysis. Authors are also grateful to Dr. I. D. Singh, Ex, Head, Plant Improvement Division, TTRI, Jorhat, Assam, Dr. Sanjay Kumar, Director, and Dr. Ram Kumar Sharma, Principal Scientist, IHBT, Palampur, Himachal Pradesh, India for providing the valuable suggestions and Mr. S. Soundararajan, Secretary, NTRF for coordination as well as Mr. Shib Nath Sinha, NTRF, Kolkata for secretarial assistance. Authors are grateful to National Tea Research Foundation, Ministry of Commerce, Kolkata, India for financial support.

